# Renewable Self-Folding Origami Constructed from Bioengineered Bacterial Cellulose

**DOI:** 10.1101/2025.08.22.671751

**Authors:** Yitong Tseo, Morgan Guempel, Cathy Hogan, Ian Hunter

## Abstract

This work presents the first genetically engineered cellulose actuator where the structural basis for actuation is directly encoded in the producing organism’s DNA. By modifying *Komagataeibacter rhaeticus* to secrete BslA protein and treating with NaOH, we create surface-activated bacterial cellulose that exhibits water contact angles 2.4*×* greater than and water retention 8.2 *×* less than unmodified cellulose. Layering this hydrophobic BslA-activated cellulose with untreated bacterial cellulose produces a biofabricated actuator driven by differential strain during dehydration. The actuation follows the Timoshenko bilayer beam equation adapted for hydrogels and can be fabricated either by shaping mature pellicles or through direct growth in 3D-printed autoclavable molds. We demonstrate two applications with a self-folding bacterial cellulose origami crane and a biomimetic bacterial cellulose gripper capable of supporting more than 360 times its own weight. This approach represents a significant advance in sustainable soft robotics through the creation of fully renewable, biodegradable, and genetically programmable actuators.

## 1 Introduction

Cellulose, the single most abundant polymer on Earth, possesses many critically useful features that have prompted humans across millennia to adopt it as the material of choice for a wide array of use cases. Cellulose is readily functionalizable, biocompatible, biodegradable, and abundantly renewable - synthesized by organisms across the full breadth of the tree of life with an estimated annual global production of 10^11^ *−* 10^12^ tons [14, 19]. Its remarkable mechanical properties derive from a hierarchical structure of semi-crystalline microfibrils composed of linear chains of *β*(1 → 4) linked D-glucose units, providing tensile strength comparable to steel when normalized by density [16].

### 1.1 Cellulose Actuators

With increasing modern demand for robotics and greater global attention towards sustainability, cellulose has emerged as a compelling foundational material for developing actuating and responsive devices from renewable sources [8, 9]. A major advantage of cellulose lies in its inherent biocompatibility, which positions cellulose-based actuators as particularly promising candidates for healthcare applications such as targeted drug delivery systems, biomedical devices, and tissue engineering scaffolds [26, 34].

The underlying mechanisms investigated by different cellulose-based actuators vary widely, ranging from localized anisotropic water-controlled swelling in 4D printed designs [21], to embedded magnetic nanoparticles within a cellulose matrix [29], to thermo-responsive and conductive polymer coatings [11, 39]. However many of these approaches require incorporating non-renewable materials into the cellulose structure (ex. lithium [27] and gold [32]), compromising the overall sustainability of the resulting actuators. In this work, we introduce a pioneering cellulose-based actuator designed from a bioengineered functional foundation. Our material relies solely upon cellulose, secreted BslA protein coating, and differential dehydration for actuation, thereby eliminating the need for bio-incompatible or non-renewable additives. Moreover, the components of our system are constructed via bacterial biofabrication, making it especially attractive for rapid, scalable, and sustainable production [35].

### 1.2 Bacterial Cellulose

Bacterial cellulose (BC) has garnered considerable attention as one of the most efficient sources of high purity (>95%) cellulose. Compared to trees and other plants which produce cellulose interlaced with lignin and hemicellulose, bacterial strains such as *Komagataeibacter rhaeticus* (*K. rhaeticus*) synthesize hydrogel biofilms (i.e., pellicles) which are composed of type I crystalline cellulose in high purity [52].

One additional benefit that BC holds over other sources (i.e. plants) is the ease of genetically engineering the source organism. After altering or adding new DNA instructions to BC-producing bacteria, the resulting cellulose material can be produced in enhanced quantities (up to a 4-fold increase) [25, 36, 41, 56], produced under resource limited conditions [3, 24, 55], co-produced with other polymers to form composite materials with new material properties [15, 46, 54], and even engineered with light-sensitive gene expression to create spatially-resolved patterns in the material [50]. Much of this ease of engineering is due to the several synthetic biology genetic toolkits made for *Komagataeibacter*, the most commonly used bacterial genus used for BC production [18, 23, 47].

Despite interest in both cellulose actuators and synthetic biology of *Komagataeibacter*, no studies were found that genetically engineered desirable actuator properties in BC or, for that matter, cellulose of any source. Here we document a novel synthetic pathway engineered into *Komagataeibacter* to induce increased hydrophobicity, and create actuators from hydrophobic-hydrophilic bilayers.

## 2 Materials and Methods

### Strains and Growth Conditions

*K. rhaeticus* iGEM cultures (ATCC, BAA-2831) were grown at 30 °C in liquid Hestrin–Schramm (HS) media (2% (w/v) glucose, 5 g/L yeast extract, 5 g/L peptone, 2.7 g/L Na_2_HPO_4_, and 1.5 g/L citric acid) [18], or on HS agar plates (1.2% (w/v) agar). Liquid cultures were maintained under either shaking or stationary conditions. Shaking cultures were supplemented with 2% (w/v) cellulase enzyme (Sigma-Aldrich, C2730) to prevent cellulose buildup and promote denser growth. Stationary cultures, used for cellulose production, were supplemented with 1% (v/v) ethanol to enhance pellicle formation [18]. Where necessary, 33 *µ*g/mL chloramphenicol (Sigma-Aldrich, DS0172) was added for plasmid selection. Although prior work used 340 *µ*g/mL [23], we observed no growth at that concentration. BslA-CBM expression was induced with 50 *µ*M N-acyl homoserine lactone (AHL) (Sigma-Aldrich, K3255).

### Inoculation of Stationary Cultures

To ensure consistent inoculation, shaking cultures were grown to turbidity, then twice centrifuged and twice washed in fresh HS media to remove cellulase as in [18]. Washed precultures were added to fresh stationary media at a 1:4 ratio and incubated at 30 °C for pellicle growth.

### Plasmid Construction and Transformation

The plasmid and component DNA parts used in this study are detailed in the Supplementary Materials publically available on GitHub. All constructs were built using the *Komagataeibacter* Tool Kit (KTK) [23]. Plasmids were assembled via Golden Gate cloning in *E. coli* Turbo (NEB), and DNA parts not already present in the KTK library were synthesized by Twist Bioscience with KTK compatible overhangs. *K. rhaeticus* electrocompetent cells were prepared and transformed by electroporation as in [18], and selected on HS agar plates containing 33 *µ*g/mL chloramphenicol.

### BC Culturing and Processing

Unless otherwise noted, BC pellicles were cultured for 10 days. After harvest, pellicles underwent two 20-minute washes in 0.1 M NaOH at 80 °C under agitation (500 rpm), each followed by a rinse in deionized water. Neutralization was achieved by submerging pellicles in dH_2_O for 1–2 hours with gentle shaking. Dried samples were deposited on glass slides or acrylic sheets and air-dried at room temperature. NaOH-treated pellicles were rehydrated in dH_2_O for 1 hour prior to downstream use. Untreated pellicles are never dehydrated, and only rinsed in dH_2_O before use.

### BslA Induction Experiments

BC pellicles were cultured in HS media supplemented with 50 *µ*M AHL to induce BslA-CBM expression. We assessed the effect of AHL induction timing on BC surface properties and mechanical behavior but found no significant differences in hydrophobicity between early and late induction, suggesting basal uninduced expression. To avoid selective bias, samples were pooled across induction schedules. All BslA-activated BC samples used in experiments and actuator construction were treated with NaOH. For the sake of brevity, this detail is not explicitly repeated throughout the text.

### Surface Wettability

Hydrophobicity was evaluated using the sessile drop technique. Contact angles of 1.00–5.00 *µ*L distilled water droplets were recorded using a custom goniometer setup: a Canon EOS 80D DSLR paired with a 100 mm macro lens (Canon EF 100mm f/2.8L Macro IS USM), backlit by an LED panel on an adjustable stage. Droplet volumes were dispensed via a precision droplet dispenser (Micro4, World Precision Instruments). Contact angles were processed using the ImageJ Contact Angle plugin [6, 44].

### Scanning Electron Microscopy

Cellulose samples were imaged using a Hitachi TM3000 tabletop SEM without the application of any conductive coating. Images were acquired under high vacuum conditions at an accelerating voltage of 15.0 kV with a working distance of 6 mm.

### Tensile Testing

Tensile testing was performed using a custom rig consisting of two FUTEK LSB200 load cells and two FUYU FSL40 Linear Guide Version 2.0 motors arranged in sequence, controlled by Tinkerforge components [22]. Sample thickness was measured using a micrometer, with care taken to secure the sample without over-tightening.

### Dehydration Analysis

Mass during dehydration was tracked for individual pellicles using a custom-built rig integrating an S-Beam load cell (FUTEK LSB200) with Tinkerforge components for control [22]. Measurements were performed in a fume hood to maintain consistent humidity.

### Differential Strain Analysis

Differential strain was analyzed by tracking the thickness over time of pellicles laid flat at 120 °C. Frame-by-frame image analysis from footage taken by a portable microscope (Jiusion 40-1000X) was conducted using ImageJ.

### Bimorph Construction

Bimorph strips were constructed by layering 5 mm *×* 20 mm strips of untreated BC on top of NaOH-treated, BslA-activated BC. To explore effects of relative thickness, untreated pellicles were cultured between 5–10 days. Actuation was induced by mounting the bimorphs above a hot plate set to 120 °C and imaged with a portable microscope (Jiusion 40-1000X).

### Bimorph Image Tracking

A custom computer vision pipeline was developed using Meta’s Segment Anything Model (SAM) [30], FFmpeg [17], and OpenCV [5]. First, the background is removed by SAM, followed by a medial axis transform to extract the bimorph skeleton [31]. Tip positions are tracked as the farthest point from the clamp, and an averaging kernel with a window size of 5 is applied. All code and data, including videos of bimorph actuation, are available at: https://github.com/YitongTseo/cellulose_origami

## 3 BslA-activated Cellulose Bioengineering and Characterization

In this section, we describe BslA-activated cellulose secreted by a genetically engineered strain of *K. rhaeticus*. BslA-activated BC fabricated in this way represents a novel biomaterial which achieves high hydrophobicity, low water logging, and rigidity even while wet. These properties establish it as a promising material for sustainable packaging, fashion, and medical treatment as well as the foundational material for our self-folding hydration based actuator.

### 3.1 Plasmid Engineering

In this work, BslA, a protein produced by *B. subtilis* which self-assembles into waterproof coating for biofilms [2], is conjugated with a cellulose binding module protein (CBM) in order to activate the surface of bacterial cellulose. Previous work by Gilmour et al. has shown that BslA-CBM constructs, when isolated from engineered *E. coli* extract and used to treat *Komagataeibacter*, significantly increases the hydrophobicity of the resultant cellulose pellicles [20]. However, direct secretion of BslA-CBM by *Komagataeibacter* was not attempted, potentially due to challenges in protein secretion engineering.

Although *Komagataeibacter* has been engineered to secrete other non-native polymers (e.g., chitin [54], hyaluronic acid [46]), only one study successfully engineered it to not only produce but secrete entire non-native proteins, doing so using a modified Type VIII secretion system from *E. coli* [23]. More specifically, they found that using the CsgG pore-forming protein along with the N22 localization tag on a protein of interest leads to optimal secretion. This suggests that CsgG, with the N22 tag, is sufficient for BslA-CBM secretion in *K. rhaeticus*.

Using the *Komagataeibacter* Tool Kit (KTK) [23], we designed a plasmid for BslA-CBM production and secretion (Figure 1). This plasmid includes the CsgG gene for protein secretion, the BslA-CBM gene with the N22 tag, and the LuxR gene for chemical inducibility via acyl homoserine lactone (AHL). In engineering *K. rhaeticus* to produce and secrete BslA-CBM, the external *E. coli* protein production and extraction steps are bypassed, thereby greatly reducing culturing overhead and streamlining the fabrication process for potential mass production.

**Fig. 1.**
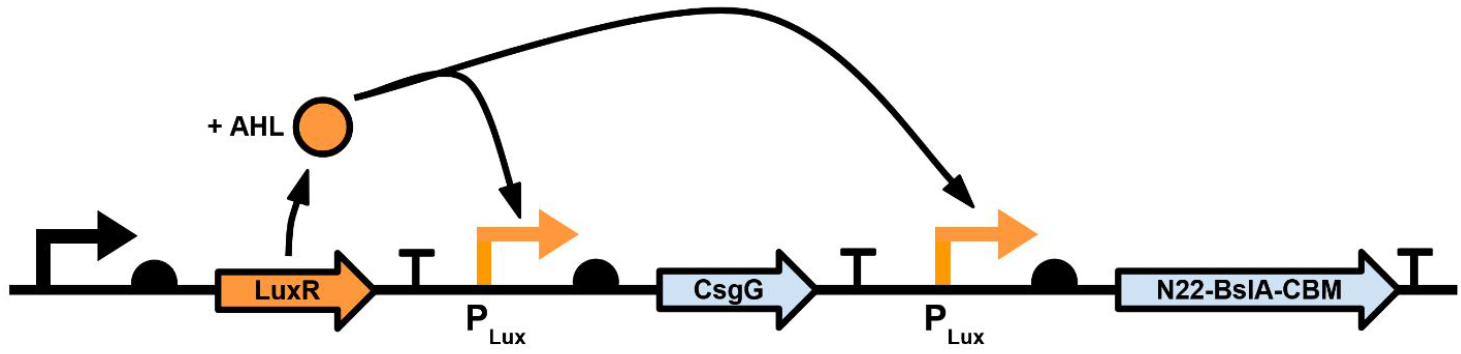
Genetic construct map for the *K. rhaeticus* BslA expression plasmid. When bound to AHL, LuxR binds to the PLux promoters and activates the production of the cellular secretion machinery protein CsgG and the secretion-tagged N22-BslA-CBM. Together this results in secretion of BslA that binds to extracellular bacterial cellulose.

### 3.2 Scanning Electron Microscopy

Figure 2a and Figure 2d show SEM images of genetically unmodified (also termed wild-type, or WT) BC and NaOH-treated BslA-activated BC pellicles after 10 days of incubation. It is qualitatively observed that untreated WT pellicles exhibit low structural rigidity and need external support to prevent folding, while NaOH-treated BslA-activated pellicles are rigid enough to self-support, even when hydrated. When viewed with SEM, a dense micro-fibrous mesh is visible, composed of cellulose fibers—each < 1 *µ*m thick (Figure 2b,c). The micro-porous topology is consistent with other SEM BC surface analyses and is known to contribute to BC’s inherent hydrophilicity [12].

**Fig. 2.**
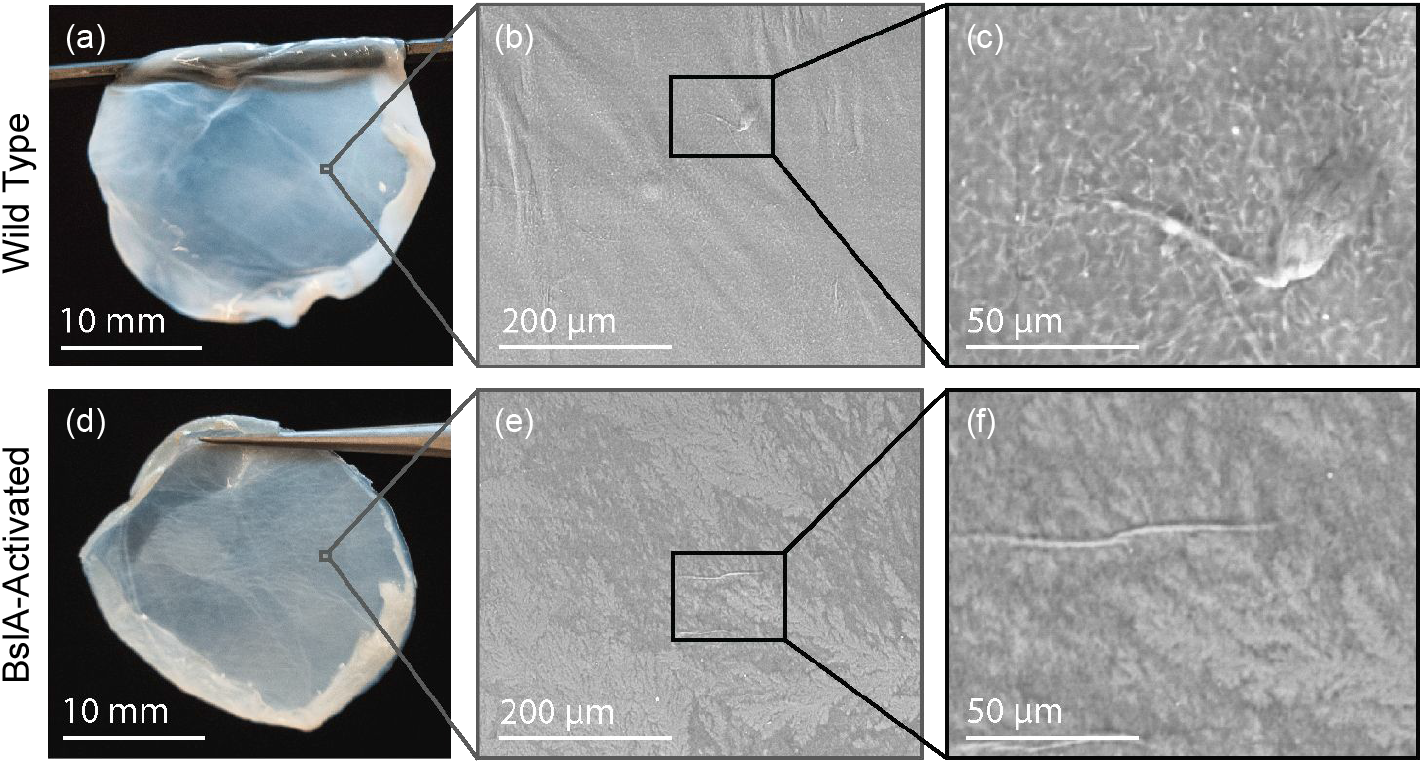
Macroscopic images and scanning electron microscopy (SEM) micrographs of BC pellicles. **(a)** Untreated WT BC. **(b)** SEM micrograph of untreated WT BC at 300x magnification. **(c)** SEM micrograph of untreated WT BC at 1500x magnification. **(d)** NaOH-treated BslA-activated BC. **(e)** SEM micrograph of BslA-activated BC at 300x magnification. **(f)** SEM micrograph of BslA-activated BC at 1500x magnification.

NaOH-treated BslA-activated BC pellicles show dark anisotropic striations not present in untreated WT samples (Figure 2e), possibly indicating a self-assembled BslA-CBM surface coat or micro-fissures from NaOH-induced hydro-lysis [20]. At higher magnification, the cellulose strands in the untreated pellicle are no longer discernible (Figure 2f). This change in nano-structure potentially accounts for the altered wettability of NaOH-treated BslA-activated BC [49].

### 3.3 Surface Wettability Characterization

Figure 3a compares contact angle distributions between NaOH-treated BslA-activated BC (Figure 3b), NaOH-treated WT BC (Figure 3c), and untreated WT BC (Figure 3d). The low contact angles of untreated WT BC (*θ*_*avg*_ = 42.8± 22.2) align with previous findings [4, 43]. It is this hydrophilic nature of untreated BC which directly poses challenges in its application in fields such as packaging and fashion [53].

**Fig. 3.**
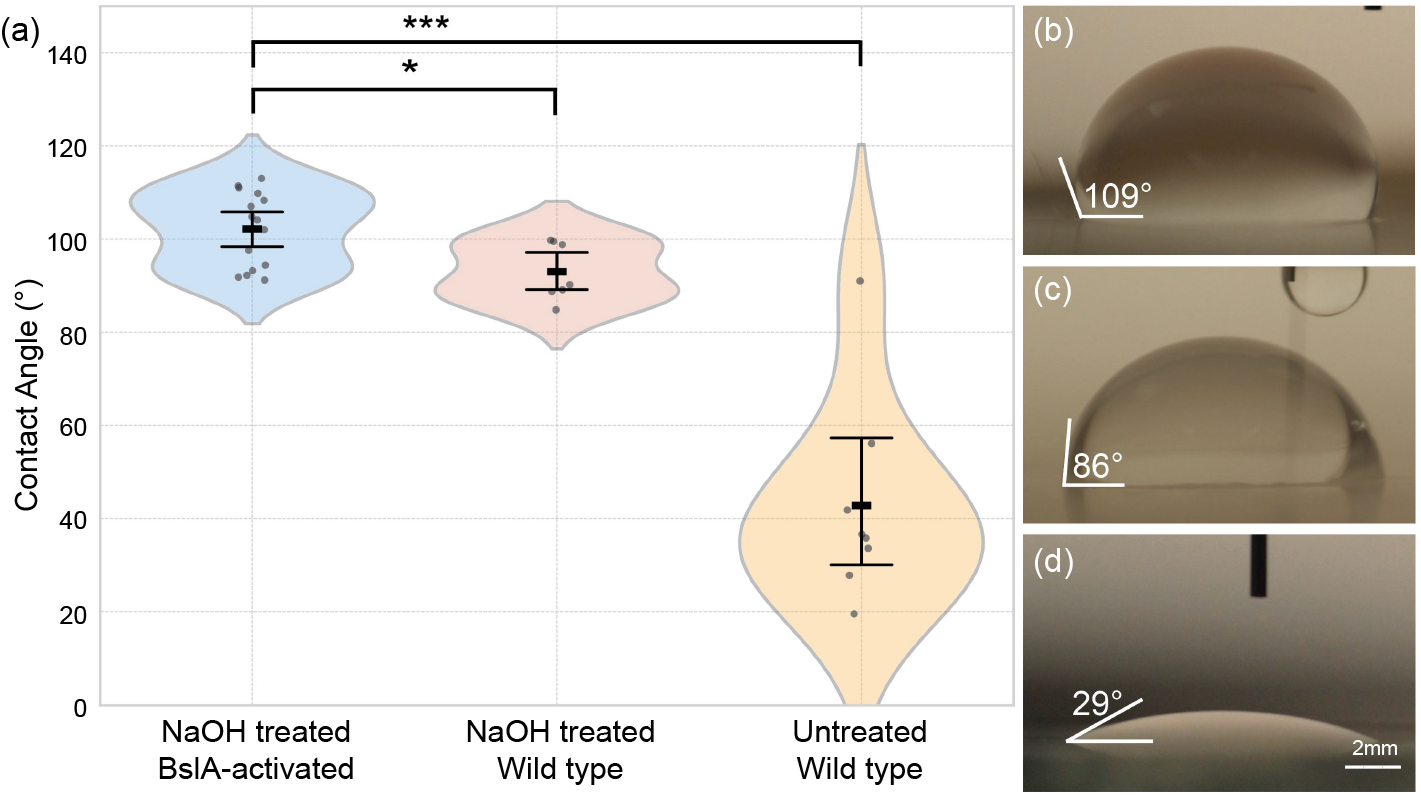
Surface wettability experiments. **(a)** Contact angle distributions across different BC treatments. **(b)** Representative contact angle formed on 0.1 M NaOH-treated BslA-activated BC. **(c)** Representative contact angle formed on NaOH-treated WT BC. **(d)** Representative contact angle formed on untreated WT BC.

From NaOH treatment, we observe on average a 2.2 *×* increase in contact angle for WT BC specimens (*θ*_*avg*_ = 93.0± 6.2), a statistically significant difference (p-value = 0.00025). BslA surface activation through our genetically modified *K. rhaeticus* strain provides an additional statistically significant 1.1*×* increase in average contact angle (*θ*_*avg*_ = 102.1± 8.0) relative to NaOH-treated WT BC (p-value = 0.0103). Our measurements for BslA-activated BC wettability align with Gilmour et al., who reported contact angles of *θ* = 105± 8.62 [20]. Notably, Gilmour et al. represent the only other known application of BslA-CBM to BC. However, their approach differs critically from ours: Gilmour et al. sources BslA-CBM from *E. coli*, requiring concurrent cultivation of both *K. rhaeticus* and *E. coli* strains and labor-intensive protein purification. Whereas our method expresses and secretes BslA-CBM directly from *K. rhaeticus*, requiring only a single culture with no purification.

We conclude that NaOH treatment combined with BslA-activation yields greater hydrophobicity than either NaOH treatment alone or no treatment. Our results seem to suggest NaOH treatment may account for most of the observed hydrophobicity increase, contradicting previous literature. Gilmour et al. found pellicles washed in NaOH (60 min, 90°C) showed no significant hydrophobicity difference relative to BslA-CBM addition, though they did not specify NaOH concentration [20]. A separate study reported BC pellicles boiled in 0.1 M NaOH for 30 min maintained tensile strain and Young’s modulus but exhibited increased swelling in SEM images; the authors, however did not specifically test wettability [38]. Another study found mild NaOH treatment (0.125 M) increased water vapor permeability [28], which may contribute to surface hydrophobicity. Regardless since surface wettability is not linearly additive [51], further experiments are needed to conclusively determine NaOH treatment’s effect on hydrophobicity.

### 3.4 Bulk Material Characterization

Differential drying rates of untreated WT BC, NaOH-treated WT BC, and NaOH-treated BslA-activated BC are shown in Figure 4a, measured at room temperature as mass over time. Notably, untreated WT BC took 11 times longer to reach equilibrium dry weight compared to treated samples (1070 min vs. 70 min and 90 min). BslA-activated BC also retained 2.6× less water than NaOH-treated WT BC, as indicated by differences between wet and dry mass. These results align with wettability data (Figure 3a) and provide additional evidence that the independent effects of BslA-activation are distinct from (though complementary to) those of NaOH treatment alone.

**Fig. 4.**
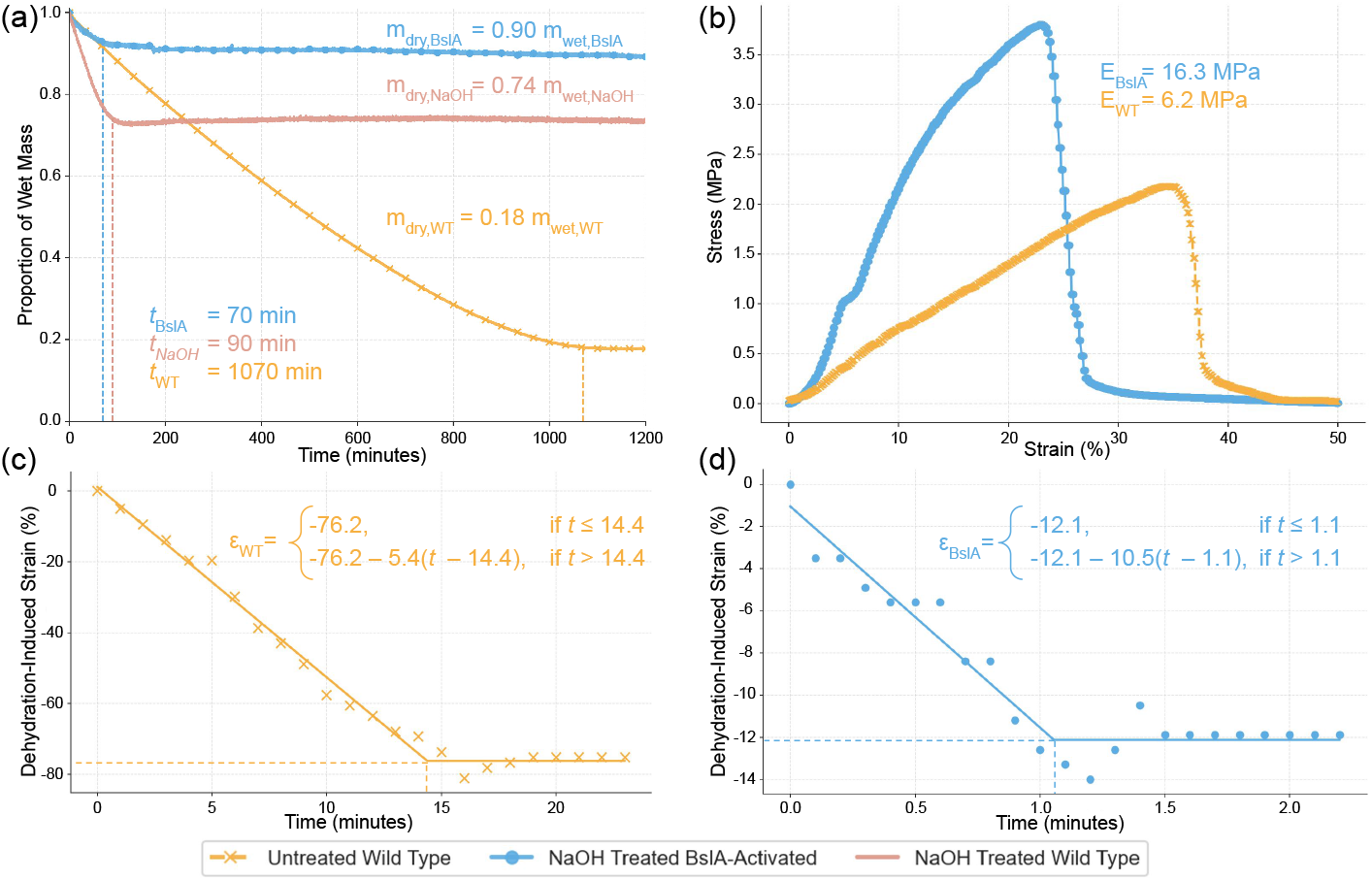
Bulk material characterization. **(a)** Mass change of hydrated BC over time as a measure of differential drying between experimental groups. **(b)** Tensile testing of hydrated BC samples. **(c)** Dehydration-induced strain as a function of time at 120 °C for untreated WT BC. **(d)** Dehydration-induced strain as a function of time at 120 °C for NaOH-treated BslA-actiated BC.

The largest differences in drying behavior and water retention were observed between untreated WT BC and NaOH-treated BslA-activated BC. Given that hydration-dependent swelling drives bimorph actuation, we focused subsequent analysis on these two materials. Dehydration-induced mechanical strain at 120 °C was tracked for WT and BslA-activated BC (Figure 4c,d); piecewise linear functions were found to fit well to the time-series data, yielding *R*^2^ of 0.946 and 0.992 respectively. Analysis revealed that BslA-activated BC experiences strain reduction close to 2x faster than untreated WT BC, as determined by the difference in initial slopes (-10.5 vs -5.4), but ceases strain reduction sooner (after 1.1 min vs 14.4 min) and achieves 6.2x less absolute strain change.

Tensile testing of hydrated samples (Figure 4b) revealed that BslA-activated BC was more rigid than WT BC, with higher peak stress and lower peak strain. Previous studies report minimal impact of mild NaOH treatment on rigidity [7, 38], though higher concentrations can weaken tensile strength [10]. In contrast, BslA incorporation appears to increase rigidity, consistent with findings by Gilmour et al. [20]. These results further support the conclusion that BslA secretion significantly alters BC material properties.

## 4 Self-Folding Bacterial Cellulose Origami

During cultivation of BC, pellicles are secreted by bacteria at the air-water interface, with each new layer forming at the top and binding to previous ones below [45]. This natural adhesion is the result of nanofibril entanglement within the BC network [1, 37]. Exploiting this evolved property, we found that hydrated untreated WT BC layered onto hydrated BslA-activated BC will bind during drying without any required adhesive. As the constructs dry, differential water retention and dehydration rates between layers generate a bending moment.

Leveraging this effect, we introduce a novel bioengineered cellulose actuator which we term **Self-Folding Bacterial Cellulose Origami**. All components of the actuator are biofabricated, biodegradable, and biocompatible making it an ideal responsive material for applications in healthcare, food processing, agriculture, sustainable fashion, and single-use packaging.

### 4.1 Characterization

Figure 5 charts the paths of dehydrating bimorph constructs and untreated WT BC unimorph controls. Bimorph constructs achieved average final horizontal displacements of 13.3 mm± 3.4 mm, significantly greater (p-value = 0.0045) than the unimorph controls’ displacement of *−* 3.2 mm± 8.8 mm (Figure 5c,d). Referencing the time-lapsed trajectories of individual bimorph and unimorph constructs during dehydration (Figure 5a,b) clarifies the behavior of each sample population: bimorphs constructs undergo gradual XY-plane-confined curving while unimorph controls exhibit stochastic spiral buckling in the full XYZ domain.

**Fig. 5.**
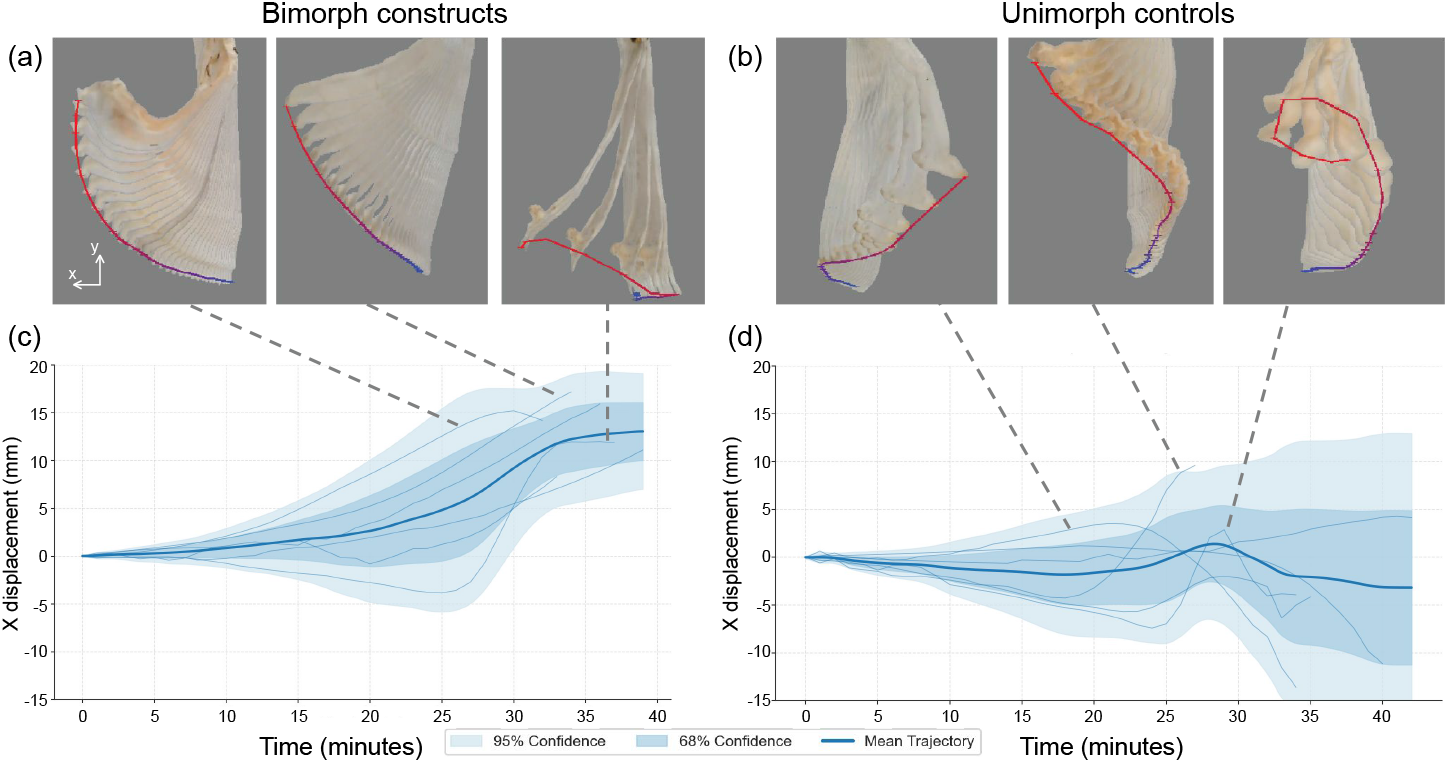
Bimorph actuation characterization. **(a)** Timelapse composites of BC origami bimorphs at 120 °C with tip migration delineated through time from blue to red. **(b)** Timelapse composites of unimorph WT BC dehydration at 120 °C. **(c)** Aggregate horizontal (X) displacement of all bimorph samples through time. All samples are positioned with the wild-type layer facing the positive X direction. **(d)** Aggregate horizontal (X) displacement of all unimorph samples.

### 4.2 Timoshenko Bimaterial Bending Model

The bending equation originally developed by Timoshenko to describe bimetal thermostats [48] has been found to adapt well to the behavior of dehydrating hydrogels [33, 40]. Building from previous work, we modify the equation to characterize self-folding BC origami,

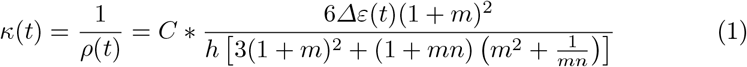

Where 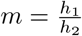 is the ratio of thicknesses of the two layers in the hydrated state, 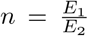 is the ratio of the Young’s moduli of the two materials as found in Figure 4b, *Δε*(*t*) = *ε*_1_(*t*) *− ε*_2_(*t*) is the differential strain of the two layers at time *t* according to experimentally derived rates (Figure 4c,d), *h* = *h*_1_ + *h*_2_ is total thicknesses of the bimorph, and *C* is the correction factor introduced by Palleau et al. [40]. To demonstrate Equation (1)‘s sensitivity, we chart the effect of untreated WT layer thickness (*h*_*WT*_) on curvature (Figure 6a).

**Fig. 6.**
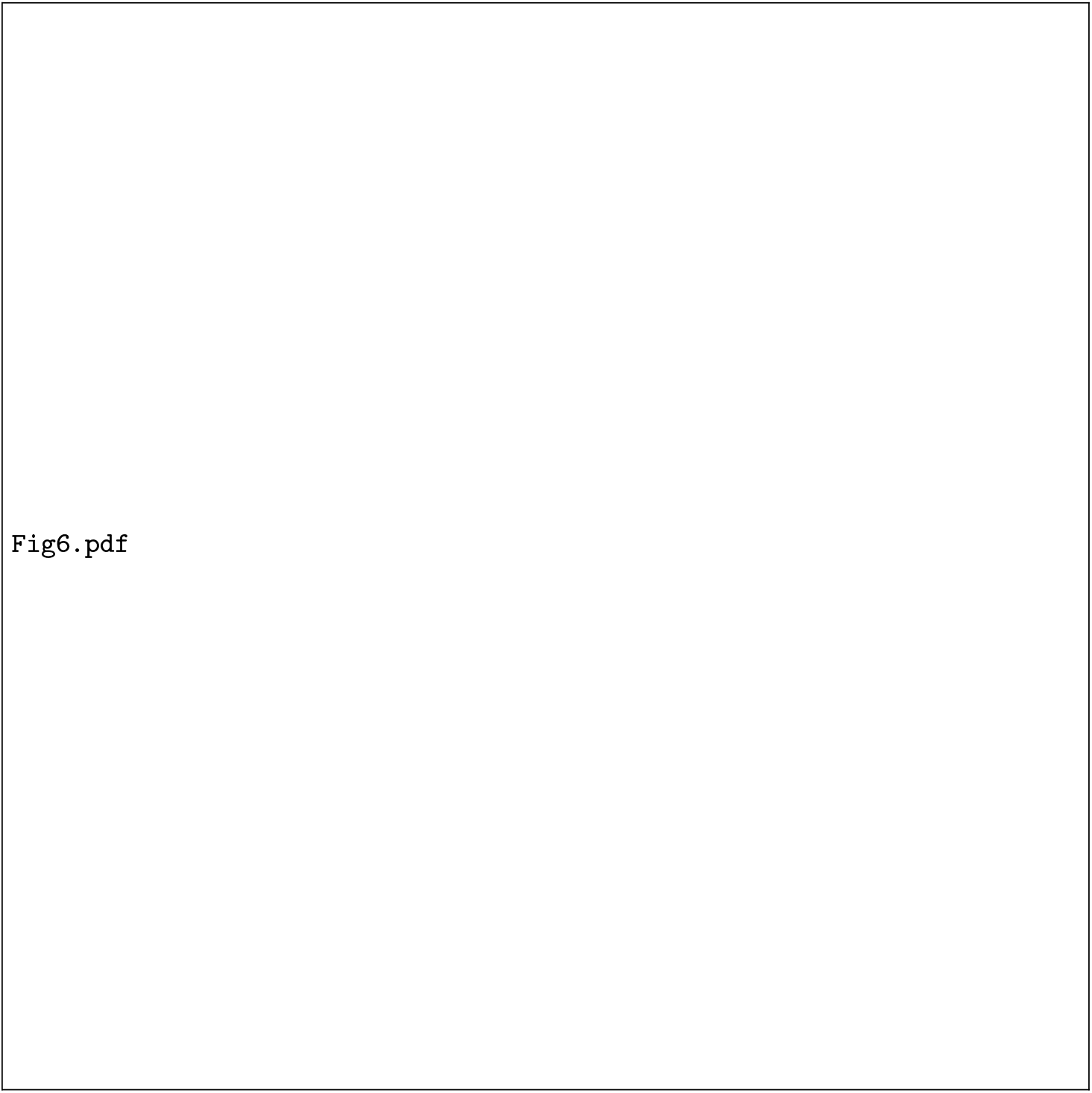
Timoshenko model adapted to self-folding BC origami. **(a)** Predicted curvature regimes at different thickness values of the untreated WT layer with the correction factor and BslA-activated thickness held constant (*C* = 1.72, *hBslA* = 0.3 mm). **(b)** Timoshenko predicted bimorph path (depicted in one-minute time intervals from pink to red) overlaid upon experimental footage time-lapsed in one-minute intervals. **(c)** Observed versus predicted horizontal trajectories plotted through time for all bimorphs.

Equation (1)‘s predictions overlaid on experimental samples at progressive time points reveal remarkable accuracy (Figure 6b). Not only does the Timoshenko model capture curvature rate and final displacement magnitude, but also explains the initial movement in the negative *x* direction. This early behavior stems from BslA-activated BC’s strain rate being 2 *×* faster than WT BC (Figure 4c,d). Only after the BslA-activated layer stabilizes does the continuing WT layer strain evolution become dominant, bending the actuator toward positive *x* at 20 min. Finally, comparing observed trajectories with model predictions across all constructed bimorphs yields a consistently low mean-squared error of 1.9 mm± 0.6 mm (Figure 6c), confirming reliable prediction of self-folding behavior across a range of *h*_*WT*_ values. For each bimorph analysis, a lag time was included for initial equilibration, and the correction factor *C* was independently optimized.

### 4.3 Self-Folding BC Origami Applications

#### Self-Folding Crane

We constructed a self-folding BC origami crane from three distinct layers: a middle untreated WT foundation layer, three BslA-activated BC segments positioned beneath the foundation to elevate the wings and neck, and a top section of BslA-activated BC to lower the crane’s head (Figure 7a). Each layer is cut from a mature BC pellicle by razor blades integrated into specially designed 3D printed molds to ensure precision. When heated to 90 °C, the crane autonomously folded within 80 minutes, resulting in a stable three-dimensional structure (Figure 7b,c). This self-folding crane demonstrates successful multi-component adhesion, precise millimeter-scale programmed actuation, and the feasibility of constructs composed of more than just two layers.

**Fig. 7.**
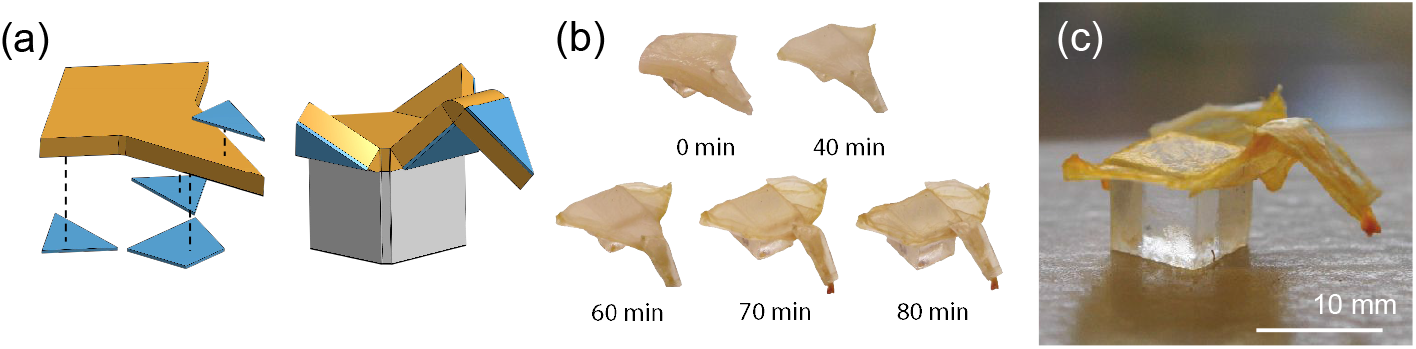
Self-folding BC origami crane. **(a)** Design of self-folding crane from layered BC. Blue pieces indicate BslA-activated BC and orange pieces indicate untreated WT BC. **(b)** Images of the BC origami crane over time. **(c)** Final structure of the crane.

#### Biomimetic Grippers

To test the carrying capacity of self-folding BC origami and to explore its application to robotics, we created a zygodactyl gripper inspired by the opposing toe structure of chameleon feet (Figure 8a). To fabricate BC with the organic gripper form, we first 3D printed molds of the shape out of autoclavable UV cured resin (Formlabs, Rigid 10K). BC pellicles were then directly grown within each autoclave-sterilized mold taking its shape (Figure 8b,c).

**Fig. 8.**
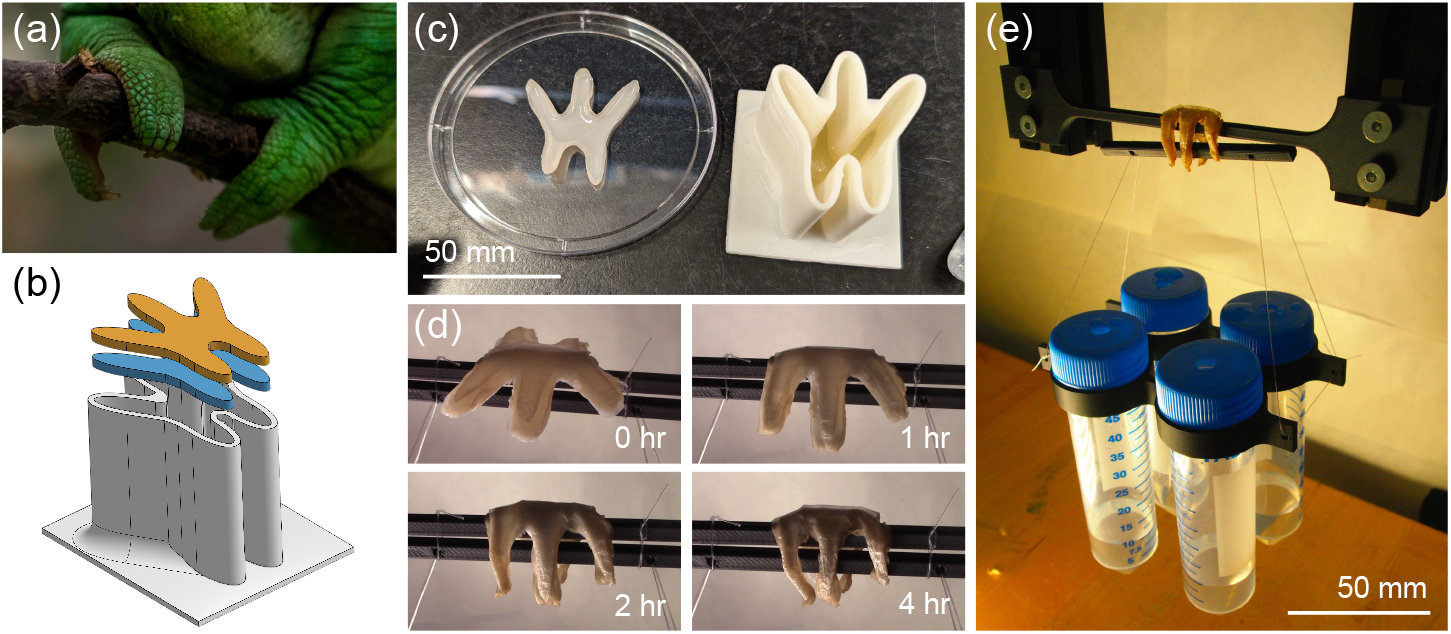
Biomimetic BC grippers. **(a)** Chameleon feet gripping a branch [13]. **(b)** Bilayer design for folding grippers. **(c)** BC pellicles grown in autoclavable molds. **(d)** Timelapse of BC gripper closing on PLA bars. **(e)** Closed gripper supporting 135g of payload.

The final assembled bilayer gripper was hung to dry at room temperature above two bars. After four hours of drying time the gripper had grasped both bars and hardened (Figure 8d). The attachments that secured the bars during the drying process were first removed, leaving only the biomimetic gripper connecting the two bars, then increasing amounts of water were added to the hanging payload (Figure 8e). The gripper failed upon the payload reaching 135 grams. This demonstrates that the gripper, which weighed only 0.37 g, was able to support more than 360× its own weight.

The successful deployment of chameleon-inspired grippers demonstrate self-folding BC origami’s potential for applications in light-weight soft robotics; with stereolithographic 3D printed molds enabling complex bio-inspired designs.

## 5 Conclusion

In this work, we present the first genetically engineered cellulose-based actuator. By modifying *K. rhaeticus* to produce and secrete BslA, a hydrophobic protein, we engineered a novel biomaterial with increased hydrophobicity, decreased water retention, and enhanced rigidity compared to unmodified BC. By layering BslA-activated and untreated WT cellulose, we then created a fully biofabricated, renewable, and biocompatible actuator that follows Timoshenko’s bimaterial model, responding predictably to dehydration-induced bending [48]. Finally, we demonstrated the actuator’s capabilities through a self-folding origami crane and a biomimetic, chameleon-inspired gripper.

BslA-activated cellulose origami offers immediate applications from auto-compressing wound dressings to sustainable self-folding food packaging and biodegradable alternatives to single-use plastics. Compared to optogenetically controlled muscle systems—biodegradable actuators that contract within seconds [42]—BC origami operates more slowly with limited cycling capability. However, BC grows more rapidly and robustly than mammalian cells, enabling cost-effective large-scale production. The material’s inherent tunability and genetic programmability present substantial room for advancement. Through further development, BC origami can serve as a platform for sustainable soft robotics, creating fully renewable, biodegradable, and genetically programmable smart materials to meet tomorrow’s environmental challenges.

## Acknowledgments

We would like to thank Inside Out, LLC for supporting this research. We extend our gratitude to Natasha Stamler for access to and instruction on the contact angle goniometer setup, Nha Nguyen for photography support, and Vineet Padia for hours of encouragement and advice. This work has received funding from NSF grant #2141064.

